# CD4^+^ T cells fuel the Duchenne cardiomyopathy

**DOI:** 10.1101/2025.10.30.684828

**Authors:** Nadine Gladow, Giuseppe Rizzo, Theresa Michel, Tobias Krammer, Antoine-Emmanuel Saliba, Stefan Frantz, Gustavo Ramos, Clement Cochain, Ulrich Hofmann

## Abstract

Duchenne muscular dystrophy (DMD) is a X-linked genetic disorder, in which cardiomyopathy represents a major cause of mortality. Although myocardial inflammation and fibrosis are hallmarks of the disease, the role of adaptive immunity in cardiac pathology remains poorly defined. Using the Mdx mouse model, we demonstrate that T and B lymphocytes accumulate in the heart and that activation occurs in heart-draining lymph nodes at an early disease stage. Mdx mice lacking adaptive immunity (Mdx-Scid) were protected from myocardial fibrosis and hypertrophic remodeling. Single-cell RNA sequencing revealed expansion of an inflammatory, matrisome-associated macrophage subset in Mdx but not Mdx-Scid hearts. Genetic deficiency or depletion of CD4^+^ T cells reduced left ventricular fibrosis and preserved systolic function. Moreover, adoptive transfer of T cells von Mdx mice induced myocardial fibrosis and dysfunction in healthy recipients. Our results identify autoreactive CD4+ T cells as key drivers of DMD-associated cardiomyopathy and suggest targeted modulation of adaptive immune responses as a potential therapeutic approach in DMD.

## Research Letter

Duchenne muscular dystrophy (DMD) is a genetic disorder which primarily manifests as progressive muscular weakness in young boys. With advances in its clinical management, heart failure has emerged as the principal cause of mortality among individuals with DMD. Cardiac magnetic resonance imaging frequently reveals signs of inflammation and histological studies reported myocardial leukocyte infiltration, which precedes decline in left ventricular function.^1^ Such findings have been interpreted as indicative of an innate immune response to myocardial injury. However, recent preclinical and clinical evidence suggests that adaptive immunity may play a causative role in the pathophysiology of DMD. Anti-dystrophin T cell responses have been documented in patients with DMD and the efficacy of corticosteroids in slowing the progression of skeletal muscle myopathy may, at least in part, depend upon the suppression of such autoreactive T cells.^2^ Of particular note, novel therapeutic strategies designed to induce expression of a functional Dmd gene product elicited autoimmune responses against dystrophin, thereby potentially compromising their therapeutic benefit.^3^

We aimed to investigate the role of adaptive immunity for development of cardiomyopathy in the Mdx mouse, a widely used model for DMD that has proven instrumental in elucidating pathological mechanisms. Compared to WT myocardium, flow cytometric analysis of hearts from three-month-old male Mdx mice on a wild-type (WT) C57BL/10 background—representing an early stage of cardiomyopathy—revealed an increased number of total leukocytes (Figure A_i_), particularly of T and B lymphocytes (Figure A_ii_). Antigen recognition induces T cell activation and proliferation in organ-draining lymph nodes. Notably, bona fide heart-draining, but not skeletal muscle-draining popliteal lymph nodes, showed significantly increased cell numbers (Figure A_iii_). Hence, at this early stage we could detect hints for lymphocyte activation. These findings suggest that an autoimmune response may precede overt cardiac pathology.

**Figure.**
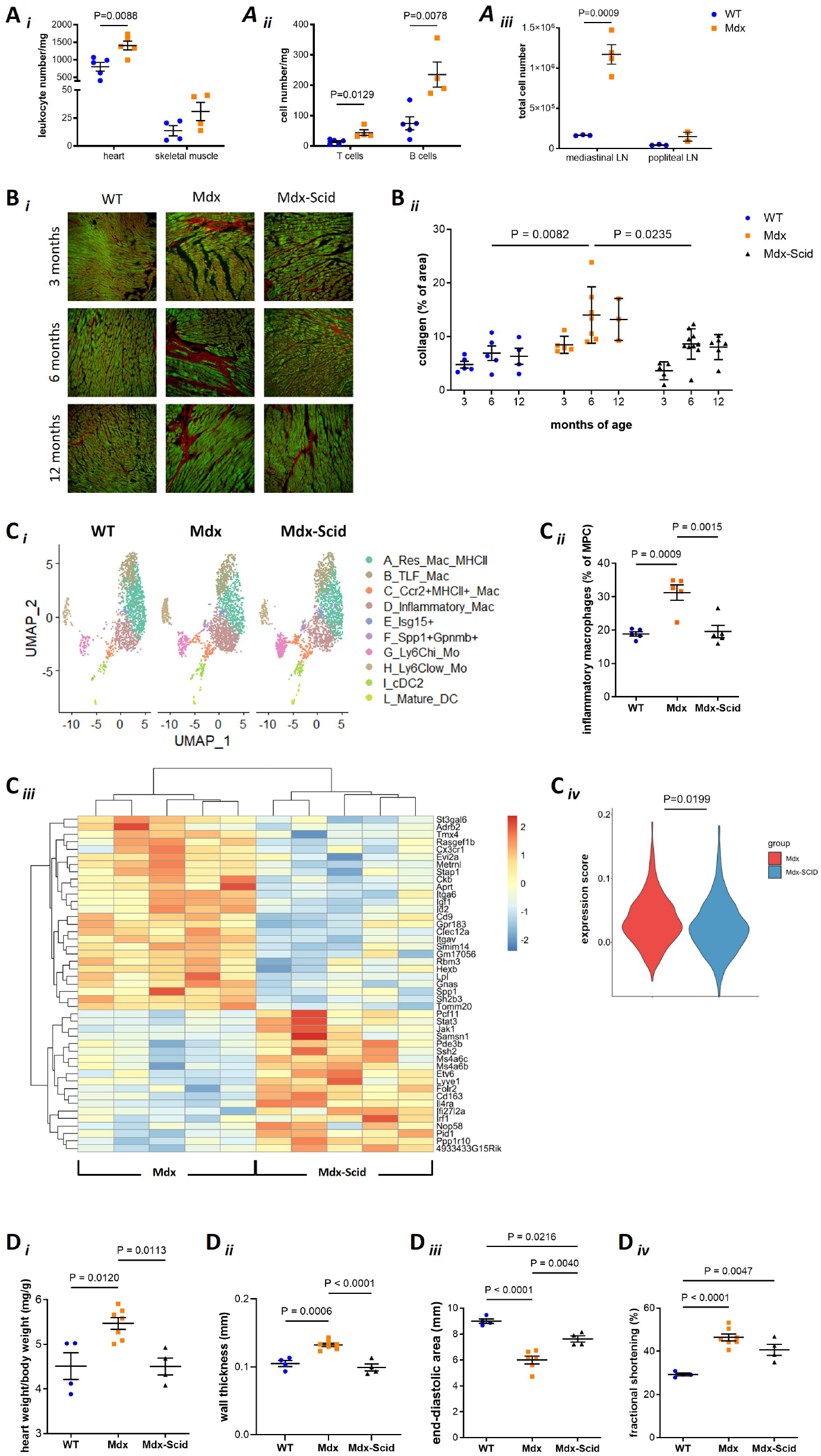
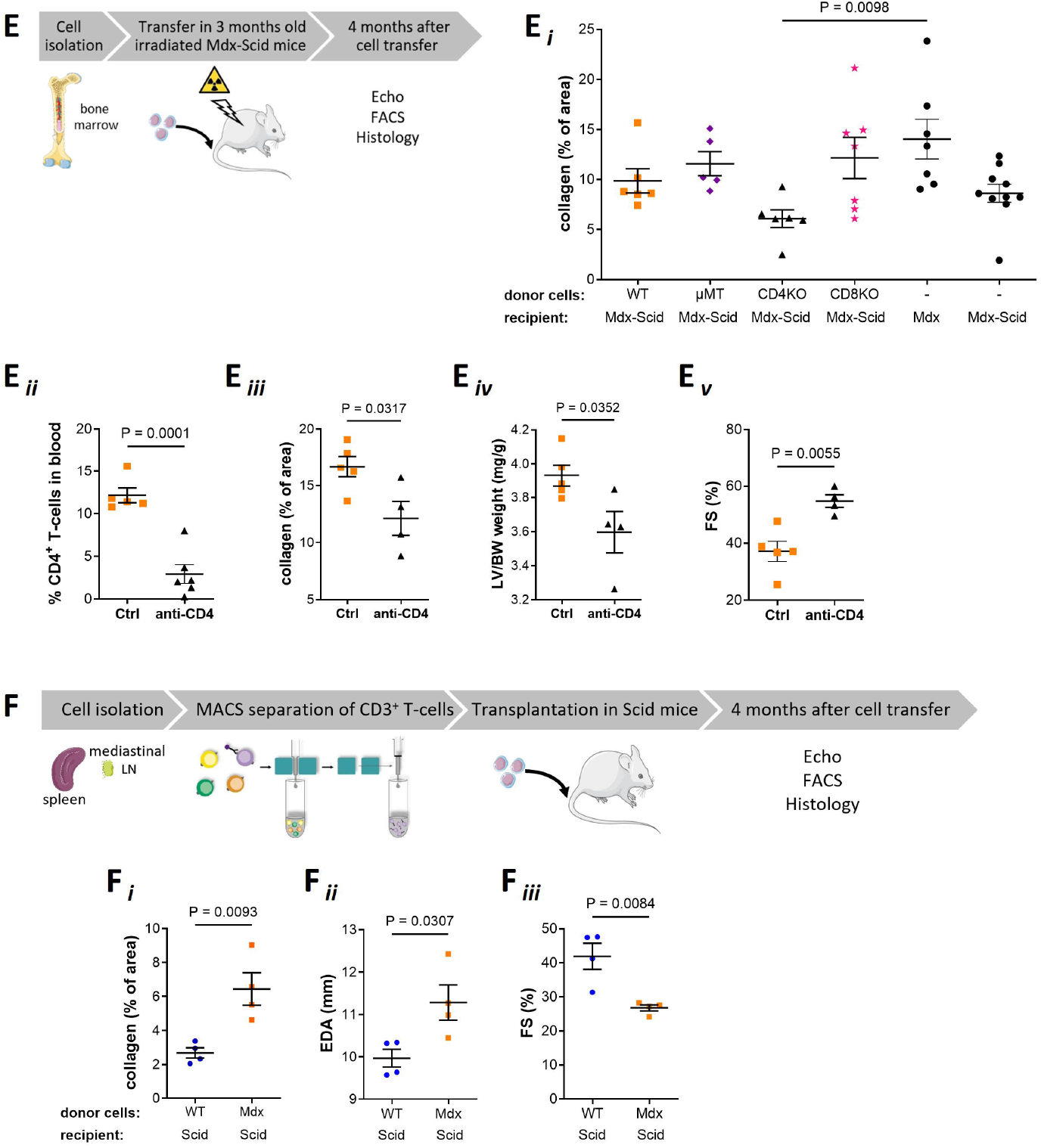
CD4^+^ T cells drive the progression of Duchenne Muscular Dystrophy cardiomyopathy. **A**, Phenotype of 3-months-old Mdx and wildtype (WT) mice. **A_i_**, CD45_+_ leukocytes in myocardium **(A_ii_**, CD3_+_ T and CD19_+_B lymphocytes), skeletal muscle and lymph nodes **(A_iii_)** analyzed by flow cytometry. **B**, Fibrosis. **B_i_**, Picrosirius red staining of collagen (red) and cardiomyocyte autofluorescence (green). **B_ii_**, Collagen quantification. **C**, Single-cell RNA sequencing of FAGS-sorted myocardial co45+ cells (1OX Genomics Chromium). **C_i_**, Uniform Manifold Approximation and Projection (UMAP) of mononuclear phagocyte (MPC) clusters. **C_ii_**, Proportion per genotype, **C_iii_** Pseudobulk analysis (using DeSeq2 package, https://www.nature.com/articles/s41467-022-35519-4) and Matrisome-associated macrophage (MAM) scores (**C_iv_** of inflammatory macrophages (cluster D) in Mdx and Mdx-SCID. **D**, Cardiac phenotype of 6-month-old mice. Organ weights **(D_i_)** and echocardiographic assessment of left ventricular (LV) structure and function (**D_ii_-D_iv_** in Mdx, WT and Mdx-Scid mice. **E**, Role of lymphocyte subsets in myocardial remodeling. **E_i_**, Fibrosis in bone marrow chimeras generated with CD4-, CD8-, or B-cell (µ MT) -deficient donors and Mdx-Scid recipients. **E_ii_**, Peripheral co4+ T cells. Myocardial fibrosis **(E_iii_)**, LV-to-body weight ratio **E_iv_**, and LV function (**E_v_**) following antibody-mediated co4+ T cell depletion (clone GK1.5 or isotype control LTF-2; i.p. every other week) in Mdx mice. **F**, Adoptive transfer of 3×10_6_ MACS-sorted T cells from WT or Mdx donors into Seid recipients. **F_i_-F_iii_**, Histological and echocardiographic analysis. Data are mean ± SEM. Normally distributed data were analyzed with unpaired t-test, one- or two-way ANOVA with Bonferroni post hoc test. MACS, Magnetic-Activated Cell Sorting; FAGS, Fluorescence Activated Cell Sorting.

To investigate the role of adaptive immunity in early cardiomyopathy development, we compared Mdx mice on either WT or severe combined immunodeficiency (Scid) background. Mdx-Scid mice are incapable of mounting adaptive immune responses. Myocardial fibrosis, an early hallmark of DMD and many other inherited cardiomyopathies, was quantified using picrosirius red staining (Figure B_i_). In three-month-old Mdx mice, but not in Mdx-Scid, we observed a significant increase in interstitial fibrosis (Figure B_ii_). Remarkably, even at 12 months of age, myocardial fibrosis remained mitigated in Mdx-Scid mice (Figure B_ii_). Given the critical role of macrophages in cardiac fibrosis, we performed single-cell RNA sequencing to gain deeper insights into macrophage states (Figure C_i_). Compared to both WT and Mdx-Scid, Mdx hearts exhibited an expansion of an inflammatory macrophage subset (Figure C_ii_). When comparing Mdx versus Mdx-Scid, genes differentially upregulated in this subset included transcripts associated with fibroblast activation (e.g., Spp1, Figure C_iii_), alongside upregulation of a gene signature characteristic of a matrisome-associated macrophage state (Figure C_iv_). To further examine the influence of adaptive immunity on cardiomyopathy progression, we studied six-month-old mice. Compared to WT, Mdx mice exhibited mild concentric left ventricular hypertrophy without systolic dysfunction on echocardiography (Figure D_i_–D_iv_). In contrast, both fibrosis and hypertrophy were significantly attenuated in Mdx vs Mdx-Scid mice.

Bone marrow transplantation experiments using CD4-, CD8-, or B cell-deficient donors and Mdx-Scid recipients revealed that Mdx-Scid mice receiving cells from CD4-deficient donors exhibited the lowest myocardial collagen content (Figure E_i_). To further corroborate the role of T cells, we depleted CD4^+^ T cells via repeated administration of anti-CD4 antibodies in Mdx mice, starting at three months of age. Anti-CD4 treatment led to significantly reduced left ventricular fibrosis in 10-month-old Mdx mice compared to those treated with isotype-control antibodies (Figure E_ii_, E_iii_). Consistent with previous findings that interferon-γ-producing CD4^+^ T cells promote myocardial fibrosis in a model of afterload-induced cardiomyopathy_4_, our results indicate that CD4^+^ T cells are essential for cardiac fibrosis in Mdx mice. Moreover, their depletion also mitigated hypertrophic remodelling and preserved systolic function in Mdx mice (Figure E_iv_,E_v_).

To further assess whether T cells from Mdx mice are sufficient to induce cardiac pathology in animals with intact dystrophin expression, we performed adoptive transfer of T cells into Scid mice. Compared to WT, T cells from Mdx donors induced myocardial fibrosis, left ventricular dilatation, and systolic dysfunction in healthy recipients (Figure F_i_-F_iii_). These findings further support the notion that cardiomyopathy in Mdx mice is driven by an autoimmune-like mechanism exerted by T cells.

Our results are in accordance with a previous report on thymic dysfunction in Mdx mice resulting in impaired central tolerance to self-antigens.^5^ From a translational perspective, our findings suggest that more specific targeting of adaptive immune responses, beyond current steroid-based therapy, merits further clinical exploration. Such approaches hold promise to synergistically delay the onset of the cardiomyopathy and enhance the efficacy of gene therapies aimed at restoring *Dmd* expression.

## Data availability

Data sets, analysis, and study materials are available from the corresponding author upon reasonable request.

## Nonstandard Abbreviations and Acronyms

DMD: Duchenne Muscular Dystrophy
Mdx: Mouse model for DMD harbouring a mutation in the Dmdgene
Scid: Severe combined immunodeficieny

## References

1. Mavrogeni S, Papavasiliou A, Spargias K, Constandoulakis P, Papadopoulos G, Karanasios E, Georgakopoulos D, Kolovou G, Demerouti E, Polymeros S, et al. Myocardial inflammation in Duchenne Muscular Dystrophy as a precipitating factor for heart failure: a prospective study. BMC Neurol. 2010;10:33. doi: 10.1186/1471-2377-10-33

2. Flanigan KM, Campbell K, Viollet L, Wang W, Gomez AM, Walker CM, Mendell JR. Anti-dystrophin T cell responses in Duchenne muscular dystrophy: prevalence and a glucocorticoid treatment effect. Hum Gene Ther. 2013;24:797–806. doi: 10.1089/hum.2013.092

3. Mendell JR, Campbell K, Rodino-Klapac L, Sahenk Z, Shilling C, Lewis S, Bowles D, Gray S, Li C, Galloway G, et al. Dystrophin immunity in Duchenne’s muscular dystrophy. N Engl J Med. 2010;363:1429–1437. doi: 10.1056/NEJMoa1000228

4. Nevers T, Salvador AM, Velazquez F, Ngwenyama N, Carrillo-Salinas FJ, Aronovitz M, Blanton RM, Alcaide P. Th1 effector T cells selectively orchestrate cardiac fibrosis in nonischemic heart failure. J Exp Med. 2017;214:3311–3329. doi: 10.1084/jem.20161791

5. Farini A, Sitzia C, Villa C, Cassani B, Tripodi L, Legato M, Belicchi M, Bella P, Lonati C, Gatti S, et al. Defective dystrophic thymus determines degenerative changes in skeletal muscle. Nat Commun. 2021;12:2099. doi: 10.1038/s41467-021-22305-x

