## Supplemental methods and materials for "CD4^+^ T cells fuel the Duchenne cardiomyopathy"

1. **Materials and Methods**
   1.1. **Animals and in vivo experiments**

Male wild-type (WT; C57BL/10ScSnJ), dystrophin-deficient (Mdx; C57BL/10ScSn-*Dmd^mdx^*/J), and immunodeficient dystrophin-deficient Mdx-Scid (B10ScSn.Cg-*Prkdc^scid^ Dmd^mdx^*/J) mice on a C57BL/10 background at an age of 2, 3, 6 or 12 month were used. Animals were housed under specific pathogen–free conditions with ad libitum access to food and water. All animal experiments were approved by the local governments (Regierung von Unterfranken) and performed according to the European Parliament Directive 2010/63/EU on the protection of animals used for scientific purposes as well as the NIH guidelines. All analyses were performed by researchers blinded to treatment and genotype.

For adoptive transfer experiments, bone marrow cells from CD4- (B6.129S2-Cd4^tm1Mak^/J), CD8- (B6.129S2-Cd8a^tm1Mak^/J), or B-cell (µMT¸ B6.129S2-Ighm^tm1Cgn^/J) -deficient donors were isolated and injected intravenously into 3-month-old Mdx-Scid recipients receiving a low-dose whole-body irradiation prior to transplantation. CD3⁺ T cells were isolated from mediastinal lymph nodes and spleens of donor mice (WT or Mdx) using magnetic cell sorting (MACS; CD3ε Micro Bead Kit, mouse, Miltenyi Biotec) according to the manufacturer’s instructions. Purity of isolated T cells was analysed by flow cytometry (FACS). Purified T cells (3x10^6^ cells per mouse) were injected intravenously into Scid mice (B6.Cg-*Prkdc^scid^*/SzJ).

In separate cohorts, 3-month-old Mdx mice received intraperitoneal injections of either anti-CD4 antibody (clone GK1.5, BioXCell) or isotype-matched IgG control antibody (clone LTF-2, BioXCell) every other week (200µg per dose) until 10 month of age.

All mice were obtained from Jackson Laboratories, except Mdx-Scid mice, which were originally provided as a breeding pair by Jackson Laboratories and subsequently maintained as an in-house colony at Charles River Laboratories.

Echocardiography was performed in spontaneously respirating animals under isoflurane (0.5-1.0 vol%) anaesthesia.

At the end of the experiments, animals were either euthanized by cervical dislocation or anesthetized for perfusion, if the organs were used for FACS analysis. Therefore, mice were injected with ketamine (100mg/kg body weight) and xylazine (10mg/kg body weight) in 100µl sterile saline. To prevent coagulation during perfusion, heparin (10 IU/g body weight in 100 µL saline) was administered intraperitoneally. Animals were perfused with 40mL phosphate-buffered saline (PBS). To label circulating leukocytes, 5 µg of BV650-conjugated anti-mouse CD45 antibody was added to the perfusion solution.

1.2 **FACS**

To analyse immune cell populations, blood, lymph nodes (LN), heart and skeletal muscle were collected. Hearts and skeletal muscle were minced and enzymatically digested with collagenase (1000U/ml; Worthington, Cat #41H12763) for 30 min. at 37°C and 750 U on a thermoshaker. For Single-cell suspensions, organes were filtered through 70µm or 100µm strainers and stained for 25 min. at 4°C with fluorochrome-conjugated antibodies against CD45 (30-F11), CD3 (145-2C11), CD11b (M1/70), CD4 (RM4-5), CD8 (53-6.7) and CD19 (6D5). All antibodies were purchased from BioLegend.

Stained cells were measured on an Attune-NxT Flow Cytometer (Thermo Fisher) and analysed with FlowJo Software (TreeStarInc, V10.4.1).

1.3 **Immunohistology**

For collagen determination, Picrosirius Red (PSR) staining was performed on 16µm cryosections, fixed in 4% formalin for 30 min. After rinsing in deionized water, staining was performed using Picrosirius Red solution (Morphohisto, Frankfurt, Germany) for 60 min. Sections were washed and dehydrated. Images were acquired using a DMi8 fluorescence microscope (Leica Microsystems).

1.4 **Single-cell RNA sequencing (scRNA-seq)**

Cardiac leukocytes from BL10, Mdx and Mdx-Scid mice were isolated and processed for scRNA-seq and cell hashing using the 10X Genomics Chromium platform. Cell Ranger version 3.1.0 was used to generate the gene expression matrix counts. Demultiplexing was performed based on Hashtag antibody signals. To identify the sample of origin, multiplets and cells with undetectable hashtag signal were excluded in Seurat v4.

Clustering analysis was performed according to standard Seurat workflow. Briefly, after removal of low-quality cells (high mitochondrial transcripts), data were normalized, scaled and Principal component analysis (PCA) was performed. Uniform Manifold Approximation and Projection (UMAP) was applied for dimensional reduction using 30 principal components (PCs). After the identification of the main cell populations, cells from the mononuclear phagocyte system (MPC) were isolated for sub-clustering analysis. Pseudobulk analysis were performed using the DeSeq2 package (https://www.nature.com/articles/s41467-022-35519-4).

1.5 **Statistical Analysis**

Data are shown as mean values ± standard errors of the mean (SEM). Normality was assessed using Shapiro-Wilk tests. Normally distributed data were analysed with unpaired t-tests, one- or two-way ANOVA with Bonferroni post hoc tests. Statistical analyses were performed with GraphPad Prism software. Data were considered to be significant with p < 0.05.

All data were obtained from independent biological replicates.
